# Zebrafish larval GPR132b differentially influences wound repair and infection control

**DOI:** 10.1101/2025.07.21.666068

**Authors:** Nayanna M. Mercado Soto, Taylor J. Schoen, John Stuntebeck, Nicholas García, Madalene Halley, Nancy P. Keller, Anna Huttenlocher

## Abstract

GPR132 (G2A), a lipid- and pH-sensing GPCR, has been implicated in both pro- and anti-inflammatory signaling, but its *in vivo* function in wound repair and infection control remains unknown. Here, we investigated the role of GPR132b, a zebrafish homolog of G2A, in regulating innate immune responses. Using CRISPR-Cas9, we generated *gpr132b* mutants and found that they exhibit enhanced wound healing following sterile injury but increased susceptibility to *Listeria monocytogenes* infection, indicating that GPR132b modulates a trade-off between wound repair and antimicrobial defense. The enhanced regrowth phenotype was associated with increased macrophage accumulation at the wound site and reduced basal expression of the pro-inflammatory cytokine *tnf-α*. Macrophage depletion suppressed the enhanced regrowth phenotype, suggesting a functional role for macrophages in GPR132b-mediated repair. Pharmacological inhibition of cyclooxygenase (COX) and 12-lipoxygenase (12-LOX) pathways mimicked the *gpr132b* mutant phenotype in wild-type larvae, indicating that GPR132b likely responds to lipid-derived signals. Together, our findings reveal that GPR132b acts as a **c**ontext-dependent regulator of innate immunity, impairing efficient tissue repair in sterile conditions while supporting pathogen resistance during infection. Our results underscore the importance of GPCR-mediated signaling in orchestrating effective responses to tissue injury and infection.

## Introduction

Wound healing requires a tightly coordinated response to promote tissue repair, eliminate pathogens, and resolve inflammation. The innate immune system plays a central role in this process by orchestrating the rapid recruitment of leukocytes such as macrophages and neutrophils, initiating antimicrobial defenses, and facilitating the resolution of inflammation (1–3). These responses are driven by a range of endogenous and exogenous chemical signals that are detected by immune cells and guide their effector functions (4). Disruption of these tightly regulated pathways can result in impaired healing, chronic inflammation, or increased susceptibility to infection (3, 5).

G-protein-coupled receptors (GPCRs) are a large family of transmembrane proteins that sense and transduce extracellular signals that modulate cellular function (4, 6). In leukocytes, GPCRs sense lipids, cytokines, chemokines and other small molecules to regulate cell cycle, development, chemotaxis, activation, and the activation and resolution of inflammation (4, 7, 8). GPR132 (also known as G2A) is a proton- and lipid-sensing GPCR expressed in most leukocytes (9–12). Its ligands include molecules released in response to injury and oxidative stress, such as lysophosphatidylcholine (LPC), sphingosylphosphorylcholine (SPC), and oxidized fatty acids like hydroxyoctadecadienoic acids (HODEs), hydroxyeicosatetraenoic acids (HETEs), epoxyeicosatrienoic acids (EETs), N-acylamides and lactate (9, 10, 13–17). Although direct binding to G2A has not been confirmed for all these ligands, substantial evidence supports receptor activation through indirect mechanisms or low-affinity interactions (14, 18, 19).

G2A has been implicated in both pro- and anti-inflammatory processes, with its functional role likely context-dependent. Several studies have shown that G2A mediates chemotaxis of T cells, natural killer (NK) cells, and macrophages in response to LPC (14, 19–23). In macrophages, G2A signaling has been associated with pro-inflammatory (M1-like) polarization and linked to pathological inflammation in diseases such as multiple sclerosis, autoimmune encephalomyelitis, and type 2 diabetes mellitus (24, 25). In addition, G2A has been shown to promote a pro-inflammatory phenotype in acute inflammation models (23). In contrast, other studies have reported that G2A promotes an anti-inflammatory (M2-like) macrophage phenotype that supports tumor metastasis (16, 26). G2A deficiency has also been associated with exacerbated inflammation in murine models of colitis, atherosclerosis, and chronic hyperalgesia, and has been shown to impair efferocytosis of apoptotic neutrophils during the resolution of sterile inflammation (18, 19, 27–32). In neutrophils, inhibition of G2A reduces the production of reactive oxygen species and TNF-α in response to bacterial sepsis (33). G2A is also expressed in non-immune cells such as peripheral sensory neurons, where it regulates pain sensitivity, and keratinocytes, which promote the expression of IL-6, IL-8, and TNF-α while suppressing cell proliferation (34, 35). These seemingly contradictory roles highlight the complex and context-dependent nature of G2A signaling, which may be shaped by ligand availability, tissue environment, and the nature of the inflammatory trigger.

Despite growing interest in G2A biology, its *in vivo* role during dynamic immune processes such as wound healing and infection remain incompletely understood. To address this gap, we used larval zebrafish as a live vertebrate model to investigate the role of GPR132b, one of the zebrafish orthologs of mammalian G2A, in the context of sterile wound and wound-associated infection (11, 17). Zebrafish larvae offer powerful advantages for studying innate immunity, including optical transparency, genetic tractability, and conservation of inflammatory signaling pathways with mammals (36–39). In addition, zebrafish larvae have been widely used to study wound healing and response to infection (38, 40–43). Using live imaging and genetic manipulation, we characterized the role of GPR132b during immune responses following sterile or infected tissues damage. We found that *gpr132b*-deficient larvae were more susceptible to infection with *Listeria monocytogenes*, displaying impaired bacterial control and tissue repair. In contrast, these mutants exhibited enhanced wound healing following sterile tail transection. These findings identify GPR132b as a context-dependent regulator of macrophage-mediated host responses and highlight a potential trade-off between inflammation-mediated defense and tissue repair.

## Results

### The GPCR GPR132b (G2A) is not required for leukocyte development in zebrafish larvae

To characterize a role for G2A in tissue repair and innate immunity, we generated a zebrafish G2A mutant. The zebrafish genome contains two *G2A* homologs, *gpr132a* and *gpr132b* (11, 44). These paralogs share 39% and 44% amino acid identity with human G2A, respectively. While both genes are retained and expressed in various adult tissues, previous studies have reported consistent expression of *gpr132b*, but not *gpr132a*, in leukocytes during the embryonic and larval stages (11, 17, 45). Prior investigations of G2A function in zebrafish larvae used morpholino (MO) antisense oligomers to transiently knock down gene expression (17, 45). Although MOs are useful for short-term gene depletion, they are not optimal for experiments extending beyond a few days due to their transient nature and potential for off-target effects. To enable long-term studies, we used CRISPR-Cas9 genome editing to generate a stable *gpr132b* null zebrafish line by inducing a 160 bp deletion in exon 2 as shown in Figure 1A (46). In addition, we generated a *gpr132a* null zebrafish line (Supplemental Figure 1A).

**Figure 1.**
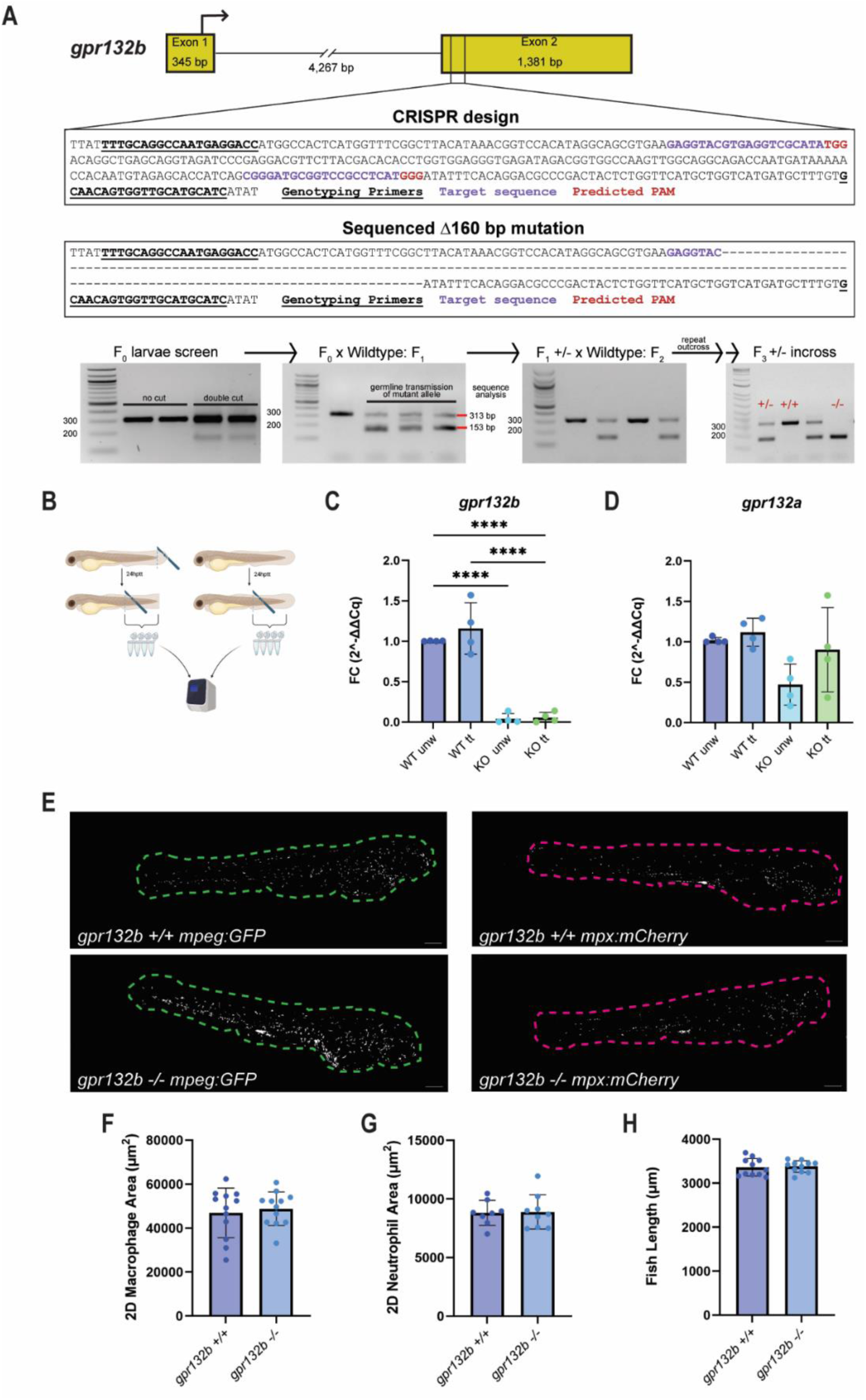
The GPCR GPR132b (G2A) is not required for leukocyte development in zebrafish larvae. (A) Schematic of *gpr132b* gene including the target region for CRISPR/Cas9 mutagenesis. Two guide RNAs (gRNAs) were used for gene editing. Genotyping primers are indicated with bold underlined text, gRNA target sequences are indicated in purple, and the PAM sequences predicted by CRISPRscan.org are indicated in red. Cutting at both gRNA target sequences generated a 160 bp deletion in exon 2 as determined by sequencing. Genotyping of F0 larvae followed by sequential outcrossing of heterozygous mutants containing the 160 bp deletion. F3 heterozygous mutants were incrossed to produce homozygous *gpr132b* mutants. (B) Schematic of RT-qPCR of wounded vs unwounded larvae. Larvae from each genotype were either transected (∼50) or left unwounded (∼50) at 3 dpf. After 24 hours, tissue from the urogenital opening to the tail fin was dissected and pooled for RNA extraction, cDNA synthesis, and qPCR. (C) RT-qPCR of *gpr132b* expression in posterior tissue at 24 hptt or 4dpf. Dots represent fold change in gene expression from pools of ∼50 larvae. Results represent 4 independent experiments. Fold change was calculated using the ddCq method. P values calculated with Two-way ANOVA. \*\*\*\**p*<0.0001. (D) RT-qPCR of *gpr132a* expression in posterior tissue at 24 hptt or 4dpf. Dots represent fold change in gene expression from pools of ∼50 larvae. Fold change was calculated using the ddCq method. P values calculated with Two-way ANOVA. Results represent 4 independent experiments. (E) Representative tile images of 3-dpf *mpeg*:GFP/*mpx*:mCherry *gpr132b* +/+ and *gpr132b* -/- larvae. Images represent maximum intensity projections of z-stacks of individual channels (scale bar is 200µm). Dashed lines outline the whole larva. (F) 2D macrophage area from whole larvae at 3 dpf. (G) 2D neutrophil area from whole larvae at 3 dpf. (H) Whole body length of 3-dpf larvae. Bars on graphs are presented as mean + standard deviation. Each dot represents data from an individual fish, and results represent data pooled from 3-4 independent experiments. n = 8-12 larvae per condition. *P*-values calculated by Student’s t-test.

To confirm the deletion of *gpr132b* and assess potential expression changes following injury, we performed RT-PCR on posterior tissue from both unwounded and tail-transected larvae. At 3 days post-fertilization (dpf), approximately 50 larvae per group were either left unmanipulated or subjected to tail transection. At 24 hours post-injury (4 dpf), the posterior region—from the urogenital opening to the tail fin—was dissected, pooled, and processed for RNA extraction and cDNA synthesis (Figure 1B). As expected, *gpr132b* transcript levels were undetectable in homozygous mutants, confirming successful knockout (Figure 1C). Additionally, we observed no significant changes in *gpr132b* expression in wild-type larvae following tail transection. Expression of *gpr132a* remained unchanged across genotypes and conditions, indicating that *gpr132a* is not transcriptionally upregulated in response to injury or the loss of *gpr132b* (Figure 1D).

Given that G2A has been implicated in regulating the cell cycle, lymphocyte proliferation, and EET-induced hematopoiesis, we investigated whether deletion of *gpr132b* affected leukocyte development in zebrafish larvae (17, 18, 47). To this end, we performed whole-larva fluorescence imaging at 3 days post-fertilization (dpf) in transgenic lines expressing GFP in macrophages (*mpeg:GFP*) and mCherry in neutrophils (*mpx:mCherry*) (Figure 1E). We quantified the total fluorescence area corresponding to each cell type across the entire larva and found no significant differences between wild-type and *gpr132b* mutant animals (Figure 1F-G). Additionally, we measured larval body length and found no significant differences between genotypes, indicating that overall growth was not affected by *gpr132b* deletion (Figure 1H). These results suggest that GPR132b is not required for normal development or maturation of neutrophils and macrophages.

### Increased susceptibility to *Listeria monocytogenes* infection in *gpr132b*-deficient larvae

GPR132 has been implicated in regulating both inflammation and host defense (23, 25, 27, 33, 34, 48). To determine whether GPR132b contributes to antibacterial immunity in zebrafish, we employed an established wound-associated infection model using *Listeria monocytogenes*, an intracellular bacterial pathogen that elicits robust innate immune responses in zebrafish larvae (40, 41).

At 3 days post-fertilization (dpf), larvae were tail transected and immediately infected at the wound site with *L. monocytogenes,* as previously described (40, 41). Infected larvae were monitored over time to assess bacterial burden and tissue damage. Compared to wild-type controls, *gpr132b* mutant larvae exhibited significantly increased bacterial burden and reduced fin tissue regrowth at 5 dpf, indicating a compromised ability to control infection and repair tissue damage associated with infection (Figure 2A-C). These findings suggest that GPR132b plays a crucial role in effective host defense against *L. monocytogenes,* and its absence results in increased susceptibility to wound-associated infection. This phenotype prompted us to investigate whether loss of GPR132b affects tissue repair in the absence of infection.

**Figure 2.**
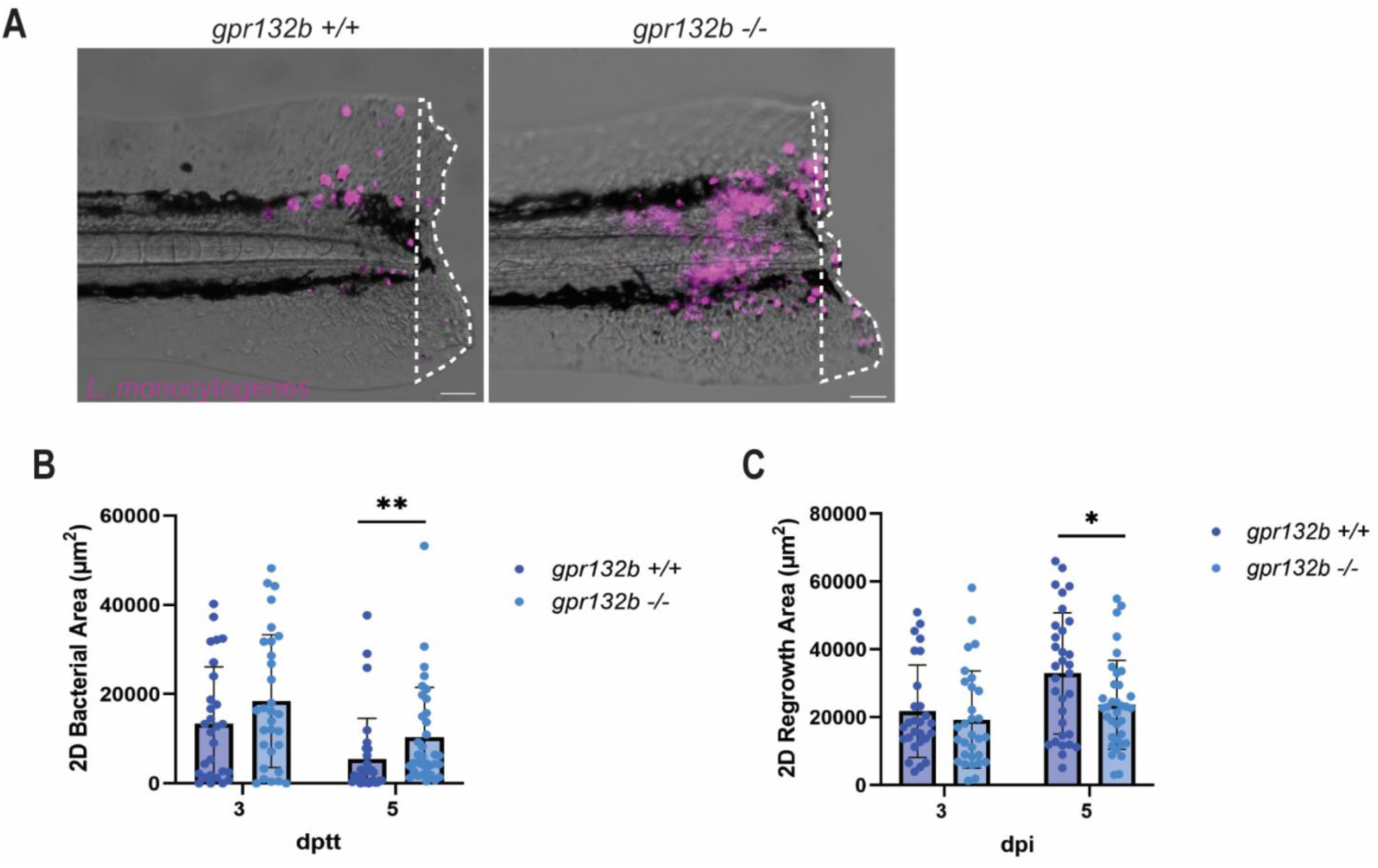
Increased susceptibility to *Listeria monocytogenes* infection in *gpr132b*-deficient larvae. (A) Representative tail fin images of *gpr132b +/+* and *gpr132b -/-* larvae infected with fluorescent *L. monocytogenes* (mCherry) at 5 dpi. Images represent maximum intensity projections of z-stacks A dashed white line outlines the regrowth area (Scale bar is 100 µm). (B) 2D bacterial fluorescent area (mCherry) at 3 and 5 dpi. (C) 2D regrowth area at 3 and 5 dpi. Bars on graphs are presented as mean + standard deviation. Each dot represents data from an individual fish, and results represent data pooled from 4 independent experiments. n = 27-35 larvae per condition. *P*-values calculated by Multiple Mann-Whitney test. \**p*<0.05, \*\**p*<0.01.

### Gpr132b-deficient larvae have improved wound healing after sterile injury

To evaluate the role of Gpr132b in tissue repair, we employed the well-established sterile tail transection model in zebrafish larvae (3, 40–42). The tail fin was amputated posterior to the notochord at 3 dpf, and tissue regrowth was assessed by brightfield imaging at 24- and 72-hours post-tail transection (hptt) (Figure 3A). We found that *gpr132b* mutant larvae exhibited a significantly larger regrowth area at both time points compared to wild-type controls, indicating enhanced wound healing in the absence of *gpr132b* (Figure 3B). To complement the stable mutant data, we generated *gpr132a/b* “crispants” by injecting guide RNAs targeting both paralogs into transgenic (*mpeg*:H2B:GFP; *mpx*:H2B:mCherry) embryos (Supplemental Figure 2A) (49). As a control for Cas9 activity, we used guides targeting *slc45a2*, a tyrosinase gene (50). At 72 hours post-wounding (hpw), *gpr132a/b* crispants exhibited enhanced tail fin regrowth compared to control crispants, although there were no differences in macrophage or neutrophil recruitment (Supplemental Figure 2B-D). To determine if *gpr132a* also contributes to tissue repair, we analyzed tail fin regrowth in larvae derived from a *gpr132a* heterozygous incross, yielding wild-type, heterozygous, and homozygous mutant siblings. At 72 hours post-tail transection, no significant differences in regrowth were observed among genotypes (Supplemental Figure 1B), suggesting that *gpr132a* is dispensable for wound healing in this context and reinforcing the idea that *gpr132b* is the functionally dominant paralog during early tissue repair in zebrafish larvae.

**Figure 3.**
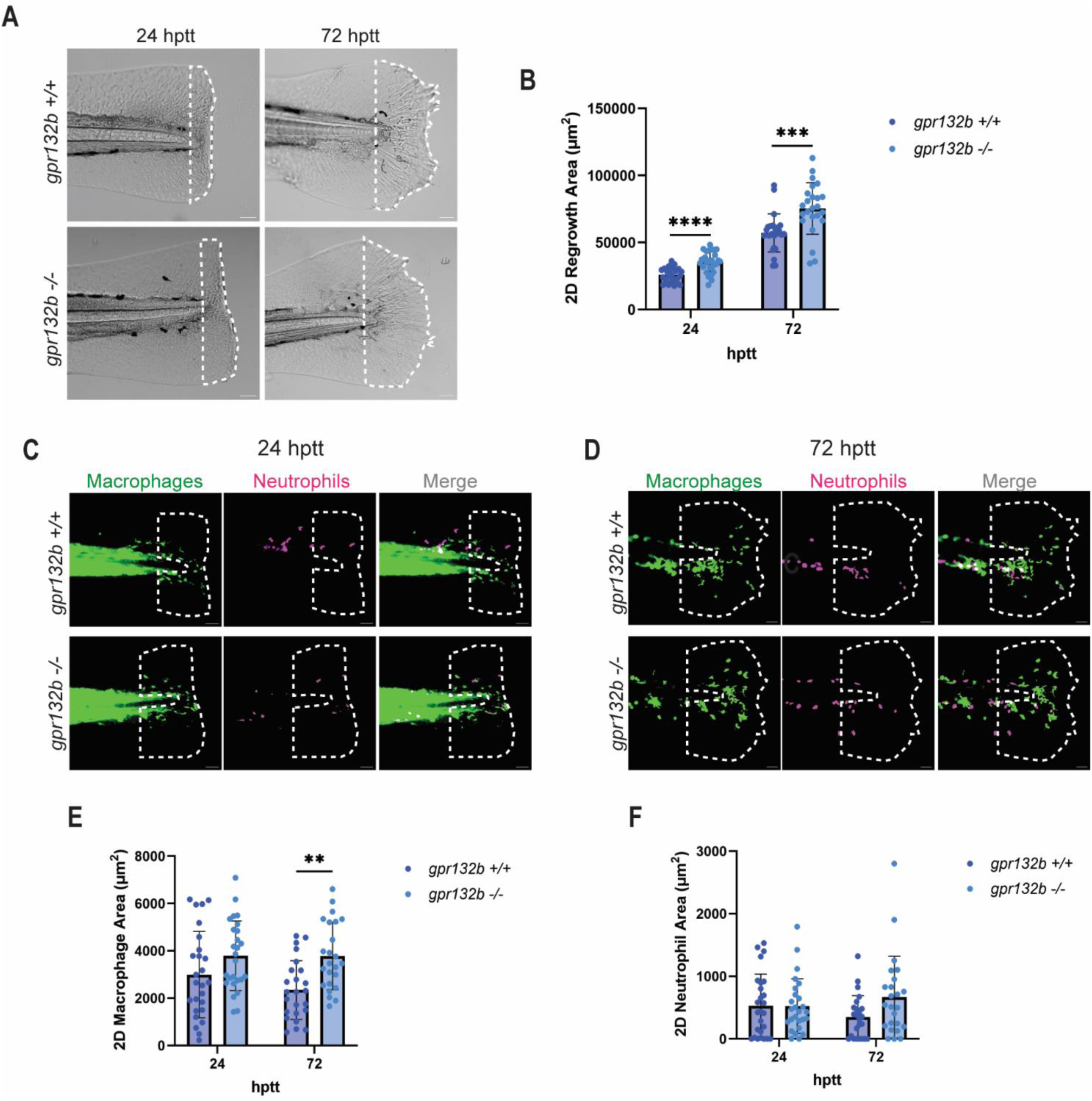
Gpr132b-deficient larvae have improved wound healing after sterile injury. (A) Representative tail fin images of *gpr132b +/+* and *gpr132b -/-* larvae at 24 and 72 hptt. A dashed white line outlines the regrowth area (Scale bar is 50 µm). (B) 2D regrowth area at 24 and 72 hptt. (C) Representative images of *gpr132b +/+* and *gpr132b -/-* larvae with fluorescent macrophages (*mpeg*:GFP) and neutrophils (*mpx*:mCherry) at 24 hptt. The dashed white line represents the region quantified, which encompasses 100μm away from the notochord. (D) Representative images of *gpr132b +/+* and *gpr132b -/-* larvae with fluorescent macrophages (*mpeg*:GFP) and neutrophils (*mpx*:mCherry) at 72 hptt. The dashed white line represents the region quantified, which encompasses 100μm away from the notochord (Scale bar is 50 µm). (E) 2D macrophage (*mpeg*:GFP) fluorescent area at 24 and 72 hptt. (F) 2D neutrophil (*mpx*:mCherry) fluorescent area at 24 and 72 hptt. Bars on graphs are presented as mean + standard deviation. Each dot represents data from an individual fish, and results represent data pooled from 3-4 independent experiments. n = 24-30 larvae per condition. *P*-values calculated by Multiple Mann-Whitney or Two-way ANOVA with Sidak’s multiple comparisons tests. \**p*<0.05, \*\**p*<0.01, \*\*\**p*<0.001, \*\*\*\**p*<0.0001.

Given these results and the known importance of leukocyte recruitment in tissue repair, we next examined immune cell dynamics following injury in the *gpr132b* mutants. Using transgenic larvae expressing fluorescent reporters in macrophages (*mpeg:GFP*) and neutrophils (*mpx:mCherry*), we quantified leukocyte recruitment at both 24 and 72 hptt (Figure 3C-D). Macrophages were the predominant leukocyte population recruited to the wound, consistent with prior studies (38, 41, 42). While macrophage recruitment was comparable between wild-type and mutant larvae at 24 hptt, *gpr132b* mutants showed a significant increase in macrophage area at 72 hptt (Figure 3E). Neutrophil recruitment remained low at both time points and did not differ significantly between genotypes, aligning with their transient role in the early phase of inflammation (Figure 3F) (3, 41, 51). These findings support the idea that macrophages play a key role in the resolution of tissue damage responses, which occurs during the later stages of wound healing (3, 41, 52).

To assess whether macrophages contribute to the enhanced regrowth phenotype in *gpr132b* mutants, we depleted macrophages using clodronate liposomes and measured fin regrowth at 24 and 72 hptt and compared wild-type and *gpr132b-*deficient larvae (Figure 4A–B)(53). Macrophage depletion significantly reduced regrowth in *gpr132b* mutants compared to wild-type siblings (Fig 4). There was very little macrophage recruitment to the wound in the presence of clodronate and no difference in macrophage recruitment between genotypes (Figure 4C). These results suggest that macrophages are necessary for the improved wound healing observed in G2A mutants. Indeed, the findings suggest that in the absence of macrophages, wound healing is impaired in the G2A mutants.

**Figure 4.**
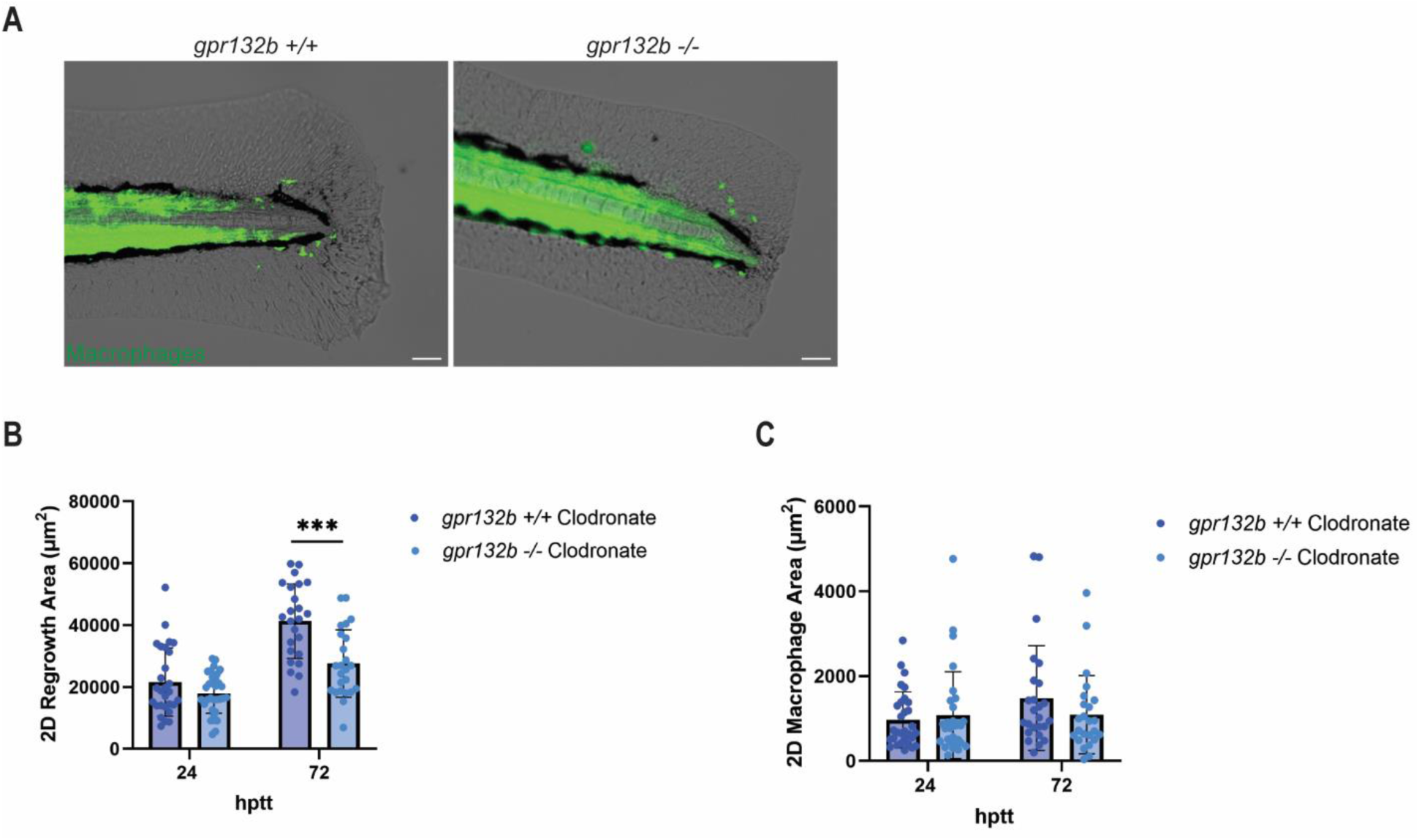
Macrophages contribute to the enhanced wound healing phenotype in *gpr132b* mutants. (A) Representative images of clodronate-injected *gpr132b +/+* and *gpr132b -/-* fixed larvae with fluorescent macrophages (*mpeg*:GFP) at 72 hptt (Scale bar is 50µm). (B) 2D regrowth area of clodronate injected *gpr132b +/+* and *gpr132b -/-* at 24 and 72 hptt. (C) 2D macrophage area (*mpeg*:GFP) of clodronate injected *gpr132b +/+* and *gpr132b -/-* at 24 and 72 hptt. Bars on graphs are presented as mean + standard deviation. Each dot represents data from an individual fish, and results represent data pooled from 3 independent experiments. n = 24-33 larvae per condition. *P*-values calculated by Multiple Mann-Whitney tests. \*\*\**p*<0.001

To determine if the inflammatory environment in wounded tissue was different in the G2A mutant, we detected inflammatory cytokine expression after wounding in control and G2A mutant fish, Our prior work showed that the predominant cell type that turns on TNF-α expression at a larval wound is macrophages (40). We measured *tnf-α* expression by RT-qPCR in posterior tissue from wounded and unwounded larvae at 24 hptt (4-dpf). While *tnf-α* levels increased modestly following injury in both genotypes, *gpr132b* mutants exhibited lower basal *tnf-α* expression relative to wild-type controls (Supplemental Figure 3A-B). These results suggest that loss of Gpr132b may modulate the inflammatory state of the tissue environment, potentially through altered macrophage activity, in a manner that supports more efficient wound repair.

### Lipid pathway inhibition enhances wound healing in zebrafish larvae

Given that several potential ligands for GPR132b are bioactive lipids, we hypothesized that pharmacological inhibition of oxygenated lipid production might similarly promote wound healing in wild-type animals (9, 10, 13–15, 17). To test this, we treated transgenic (*mpeg*:GFP/*mpx*:mCherry) wild-type and *gpr132b* mutant larvae with a combination of the pan-COX (cyclooxygenase) inhibitor indomethacin and the 12-LOX (12-lipoxygenase) inhibitor ML355, at concentrations previously validated for use in zebrafish larvae (54–56). Drug treatment was initiated at 3 days post-fertilization (dpf), three hours before tail transection, as shown in Figure 5A. Following injury, larvae were maintained in fresh drug-containing E3 media and imaged at 24 hours post-tail transection (hptt). Control groups were treated with vehicle (DMSO).

**Figure 5.**
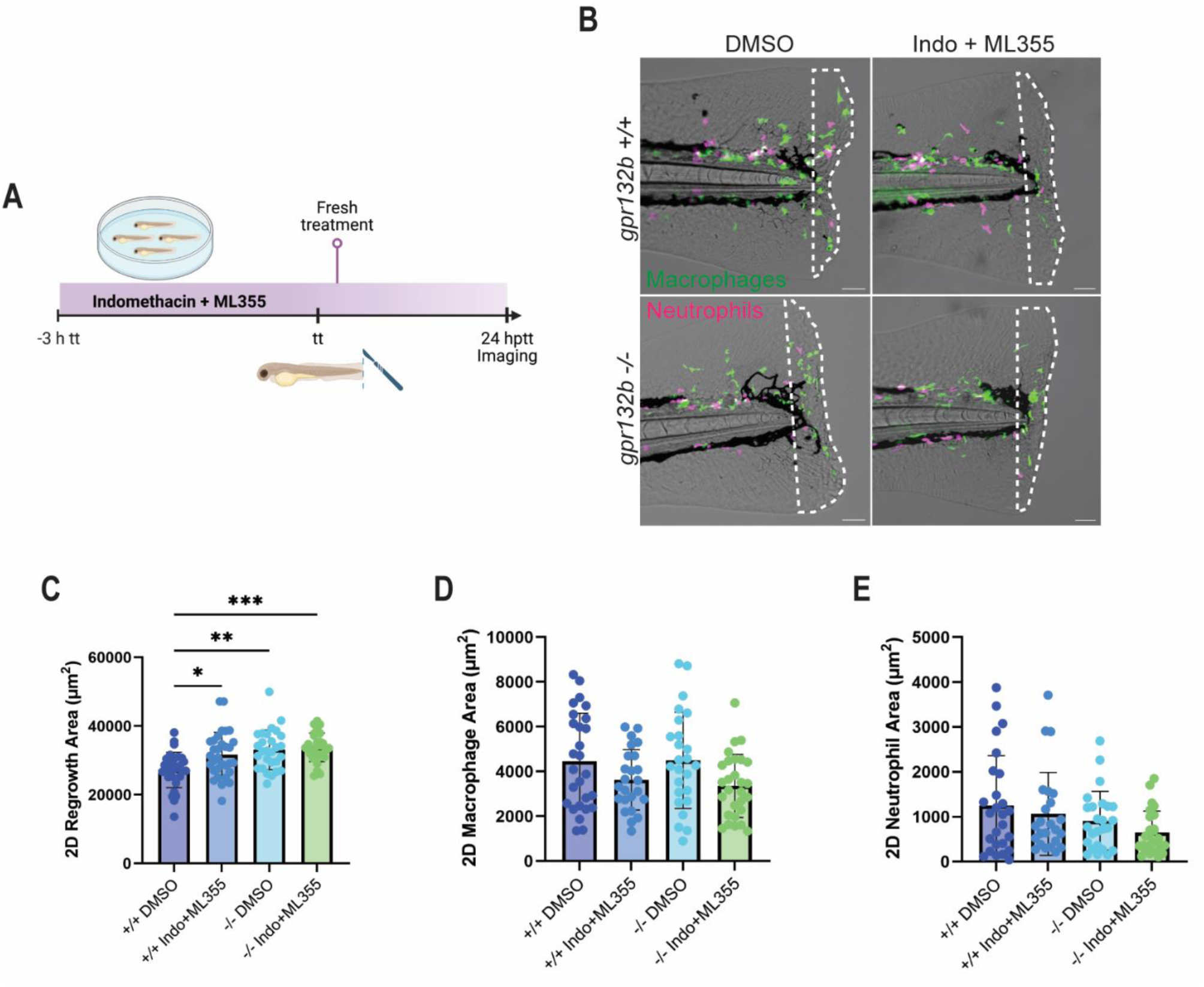
Lipid pathway inhibition enhances wound healing in zebrafish larvae. (A) Schematic of inhibitor treatment. Larvae were treated with a combination of Indomethacin (10µM) and ML355 (10µM) 3 hours before tail transection, placed in fresh solution after wounding, and imaged 24 hptt. Control larvae were treated with DMSO. (B) Representative images of treated vs untreated *gpr132b +/+* and *gpr132b -/-* larvae with fluorescent macrophages (*mpeg*:GFP) and neutrophils (*mpx*:mCherry) at 24 hptt. The dashed white line outlines the regrowth area (Scale bar is 50µm). (C) 2D regrowth area of treated and untreated larvae at 24 hptt. (D) 2D macrophage (*mpeg*:GFP) fluorescent area of treated and untreated larvae at 24 hptt. (E) 2D neutrophil (*mpx*:mCherry) fluorescent area of treated and untreated larvae at 24 hptt. Bars on graphs are presented as mean + standard deviation. Each dot represents data from an individual fish, and results represent data pooled from 3-4 independent experiments. n = 28-32 larvae per condition. *P*-values calculated by Two-way ANOVA with Tukey’s multiple comparisons tests. \**p*<0.05, \*\**p*<0.01, \*\*\**p*<0.001.

Wild-type larvae treated with the COX/12-LOX inhibitor combination exhibited significantly improved tail fin regrowth compared to DMSO-treated controls, reaching levels comparable to untreated *gpr132b* mutants (Figure 4B-C). In contrast, *gpr132b* mutant larvae did not show further enhancement in regrowth following treatment, suggesting that the effect of lipid pathway inhibition is mediated through, or converges with, GPR132b signaling (Figure 5C). Interestingly, this enhanced regrowth occurred without significant changes in neutrophil or macrophage recruitment, indicating that the effect is unlikely to be due to alterations in gross leukocyte accumulation at the wound site (Figure 5D-E). These findings support a role for lipid-derived signals in mediating Gpr132b effects during tissue repair and suggest that perturbing lipid synthesis pathways can influence wound repair outcomes in a manner that overlaps with *gpr132b* deficiency.

## Discussion

Tissue injury activates a tightly regulated inflammatory response that coordinates both repair and defense (1, 3). While this response is critical for restoring tissue integrity and eliminating potential pathogens, it must be precisely controlled to avoid excessive damage or impaired healing. G-protein coupled receptors (GPCRs) are heavily involved in cell-cell signaling in these contexts (4). Here, we examined the *in vivo* function of the GPCR GPR132b (G2A) in sterile injury and wound-associated infection. We found that *gpr132b*-deficient larvae exhibited increased susceptibility to *L. monocytogenes* infection but enhanced wound healing under sterile conditions, suggesting that GPR132b modulates innate immunity based on environmental context.

Loss of GPR132b increased bacterial burden and decreased wound regrowth, suggesting GPR132b is important for host defense during *L. monocytogenes* infection. These data are in line with previous work, which shows that G2A plays a protective role during sepsis in both *in vitro* and murine models (48). Specifically, G2A deficiency results in aberrant activation of Kupffer cells and peritoneal macrophages during bacterial sepsis, leading to reduced bactericidal activity, decreased ROS production, and increased IL-10 expression, a key anti-inflammatory cytokine (48). We have previously shown that wounds infected with *L. monocytogenes* are highly inflammatory and recruit more neutrophils and pro-inflammatory TNF-α+ macrophages than sterile injury (41). This hyperinflammation is in part due to persistent inflammasome activation mediated through IL-1β, and blocking these signaling pathways in the early stages of infection can improve healing outcomes independent of bacterial burden (40). The observation that G2A activation does not affect IL-1β expression and the poor tissue regrowth observed in our *gpr132b* mutants upon infection suggests that G2A signaling is unlikely to converge on inflammasome activation in this context (23). Therefore, we posit that the overwhelming bacterial burden due to the decreased bacterial killing capacity of G2A-deficient macrophages, as observed in other models, is responsible for defective wound healing (48). Future investigations of how G2A influences the inflammatory state of *L. monocytogenes*-infected wounds will clarify this point.

G2A activation by its ligands 9-HODE and LPC is known to induce immune cell chemotaxis (14, 20, 21). However, we observed that loss of GPR132 instead increased macrophage recruitment to the wound site. Additionally, we saw no effect of exogenous 9-HODE on early macrophage recruitment to a sterile injury (data not shown) nor in wound healing. Our data suggest that G2A signaling plays an inhibitory role in macrophage recruitment to the wound site. One possibility for this discrepancy is that G2A-mediated macrophage recruitment to this injury type does not rely on these lipid signals.

We therefore investigated how lipid signaling contributes to GPR132b-mediated wound healing by treating larvae with indomethacin (COX inhibitor) and ML355 (12-LOX inhibitor), which target enzymes involved in the production of oxygenated lipids. A limitation of our study design is that both pathways were inhibited simultaneously, and thus, we cannot determine which specific ligands or enzymes mediate the observed effect, nor can we rule out contributions from downstream metabolites. However, indomethacin has been previously reported to enhance normal wound healing (57). In line with this observation, combined COX/LOX inhibition enhanced fin regrowth in wild-type larvae, mimicking the *gpr132b* mutant phenotype. In contrast, mutants showed no additional improvement, suggesting that the effect requires or converges on GPR132b signaling. In this model, COX/LOX signaling through G2A, whether directly or indirectly, could promote a shift towards pro-inflammatory behavior of macrophages that delays wound healing in wild-type animals. This model also supports our findings that clodronate-mediated macrophage depletion reduced wound regrowth in *gpr132b*-deficient mutants compared to wild-type clodronate-treated larvae. We also found reduced TNF-a expression in the wound in the absence of *gpr132b*. Although the cellular sources of TNF-α were not confirmed, these findings suggest that *gpr132b* loss promotes a shift toward a more pro-resolving inflammatory state, which may support enhanced tissue repair, potentially through macrophage reprogramming. In the context of infection, COX inhibition has been shown to render mice more susceptible to *L. monocytogenes* and the fungal pathogen *Aspergillus fumigatus* (54, 58). Future studies will be necessary to determine whether these inhibitors also increase susceptibility to *L. monocytogenes* in our model, or whether *gpr132b*-deficient larvae are more broadly vulnerable to other pathogens.

Although our study focused on immune responses, GPR132b may also function in non-immune cells. Interestingly, wound healing was impaired in the *gpr132b*-deficient larvae depleted of macrophages, suggesting that G2A may play pro-healing roles in other cell types in the wound microenvironment. In mammals, G2A is expressed in keratinocytes and responds to reactive oxygen species to modulate proliferation and cytokine production (34). Since our transcriptional data were not cell-type specific, epithelial cells may also contribute to the altered wound environment in *gpr132b* mutants. These findings underscore the complexity of tissue repair and suggest that GPR132b may coordinate inflammatory responses across multiple cell types.

In summary, GPR132b acts as a context-dependent regulator of innate immunity in zebrafish, impairing tissue repair under sterile conditions while promoting host defense during infection. Through genetic and pharmacological approaches, we show that GPR132b influences wound repair, macrophage recruitment, and cytokine expression. These findings underscore the crucial role of GPCR signaling in maintaining a balance between wound repair and antimicrobial protection. Future studies should aim to define cell-specific roles, identify endogenous ligands, and assess the conservation of these mechanisms in mammals.

## Materials and Methods

### Ethics Statement

The Institutional Animal Care and Use Committee (IACUC) at the University of Wisconsin-Madison College of Agricultural and Life Sciences (CALS) approved the use of zebrafish in this research. The animal care and use protocol M005405-A02 adheres to the guidelines established by the federal Health Research Extension Act and the Public Health Service Policy on the Humane Care and Use of Laboratory Animals, led by the National Institutes of Health (NIH) Office of Laboratory Animal Welfare (OLAW).

### Zebrafish Husbandry and Maintenance

Adult zebrafish were maintained under a light/darkness cycle of 14/10 hours, as previously described (38). For experiments, embryos were collected, screened, and maintained at 28.5°C. All zebrafish lines utilized in this study are listed in Supplemental Table 1.

### Generation of GPR132 zebrafish mutant

All primers utilized in this study are listed in Supplemental Table 2. The protocol for generating *gpr132b* and *gpr132a* mutants was primarily based on previously outlined methods (46), with specific details provided in Figure 1A and Supplemental Figure 1A. The https://CRISPRscan.org website was employed to predict two gRNAs for each gene, which were used as a pair, ensuring that each had a CRISPRscan score of 58 or higher. gRNA DNA template synthesis was conducted using the GoTaq Green Master Mix (Promega) along with a universal primer (46) and a gRNA primer specific to the target gene. In vitro transcription of the gRNAs and subsequent DNase treatment were performed using the HiScribe T7 RNA synthesis kit (New England Biolabs) per the manufacturer’s instructions and employing 150 ng of DNA template. The purified gRNAs were stored at -80℃.

For each gene, an injection mix was prepared containing two gRNAs (final concentration of 67 ng/µl each) and purified Cas9 protein (final concentration of 200 ng/µl, PNA Bio CP01). Wild-type embryos (NHRG-1) were microinjected with 3 nl of the injection mix at the 1-cell stage. At 3 dpf, genomic DNA was extracted from the injected larvae and subjected to genotyping PCR to evaluate the efficiency of the gRNAs. The results of the genotyping are shown in Figure 1A. Adult F0 fish were outcrossed to wild-type, and progeny were screened through genotyping PCR to identify the desired mutations. Fins from heterozygous F1 larvae were collected for genomic DNA extraction and sequencing. For sequencing, the target region was PCR amplified and cloned into a TOPO TA (Thermo Fisher) vector following the manufacturer’s instructions. The plasmids were then transformed into *E. coli*, purified using the NucleoSpin Mini-Prep kit (Macherey-Nagel), and sequenced at Functional Biosciences (Madison, WI). Transgenic fluorescent lines were generated by outcrossing to heterozygous mutants and genotyping for homozygous wild-type and mutants. For crispant experiments, embryos injected with gRNAs were used at 3dpf. For some parts of this study, heterozygous were in-crossed and genotyped after experiments.

### Zebrafish tail wounding and infection

3-dpf larvae were anesthetized in 5 mL of E3 media containing 0.2mg/mL Tricaine (ethyl 3-aminobenzoate, Sigma) in a 60mm tissue culture-treated dish (Corning). Caudal fin tissue distal to the notochord was transected using a surgical blade (Feather No. 10) as previously described (40). After injury, larvae were placed on a new dish and rinsed with E3. Larvae were kept at 28.5°C until imaging or fixation.

For wound infection experiments, zebrafish larvae were immersed in 5 mL of E3 medium containing Tricaine and 100 μL of bacterial suspension and injured. For uninfected controls, 100 μL of sterile PBS was added instead of bacterial suspension. Following transection, larvae were transferred to tissue culture–treated dishes and incubated for 1 hour on a horizontal orbital shaker (75 to 100 rpm). Larvae were then rinsed with E3 medium and maintained at 28.5°C until fixation or imaging.

### Fixation

Zebrafish larvae were fixed overnight at 4°C in 1.5% formaldehyde (Polysciences, Warrington, PA) prepared in a solution containing 0.1 M Pipes (Sigma-Aldrich), 1.0 mM MgSO₄ (Sigma-Aldrich), and 2 mM EGTA (Sigma-Aldrich) (40). After fixation, the samples were washed and stored in PBS at 4°C until imaging.

### RNA Extraction and RT-qPCR

Tissue from 50-55 larvae was collected in ice-cold PBS as described in Figure 3A. Tissue was pooled, and RNA was extracted using RNAeasy Mini Kit (QIAGEN). cDNA was then synthesized using SuperScript III RT and oligo-dT (Invitrogen). Using cDNA as a template, quantitative PCR (qPCR) was performed using FastStart Essential DNA (Roche) in a LightCycler96 (Roche). Fold changes were calculated, normalized to *rps11* using the ΔΔCq method (59). Primers used are listed in Supplementary Table 2.

### Listeria monocytogenes inoculum preparation

Inoculum for *L. monocytogenes* infections was prepared as described previously (40). A streak plate of *L. monocytogenes* strain 10403S WT strain-mCherry was generated from frozen stock and incubated at 37°C (40, 60). A single colony was inoculated into 1 mL of brain–heart infusion (BHI) medium (Becton, Dickinson and Company, Sparks, MD) and cultured statically overnight at 30°C to reach stationary phase. The overnight culture was diluted 1:4 in fresh BHI and incubated for 1.5 to 2 hours to reach mid-logarithmic growth phase (OD₆₀₀ ≈ 0.6–0.8). One milliliter of the mid-logarithmic culture was centrifuged, washed three times in sterile phosphate-buffered saline (PBS), and resuspended in 100 μL PBS.

### Pharmaceutical inhibition of COX and 12-LOX

Three hours before tail transection, larvae were exposed to 10μM of pan-COX inhibitor indomethacin (Sigma-Aldrich) and 10μM of the 12-LOX inhibitor ML355 (Cayman Chemical) in E3 (Figure 4A). Both drugs have been used in zebrafish larvae at the used concentrations (54, 61). 1000X stock solutions were made in DMSO, and 0.1% DMSO was used as a vehicle control. Larvae were removed from the treatment during the tail transection and placed in E3 with both inhibitors until imaging.

### Clodronate liposome injection

Larvae were microinjected at 2-dpf into the caudal vein plexus with 2nL of clodronate or PBS liposomes (Liposoma) with 0.1% phenol red for visualization, as previously described (53).

### Live imaging acquisition

For live imaging acquisition, larvae were anesthetized and placed on their side in a zWEDGI device. Alternatively, larvae were embedded flat on their side in a 35mm glass-bottom dish (CellVis) by adding 1-2% low-melting-point agarose on top. For fixed larvae, only their tail fins were imaged after cutting them. Single plane and Z-series images (25μm) were acquired on a Zeiss Zoomscope (EMS3/SyCoP3; Zeiss, Oberkochen, Germany; Plan-NeoFluar Z objective; 112X magnification (0.7 μm resolution, 2.1 mm field of view, 9 μm depth of field) and Zen software (Zeiss). For whole fish imaging, Z-series images (5µm slices, 10x magnification) were acquired on a spinning disk confocal microscope (CSU-X; Yokogawa) with a confocal scan head on a Zeiss Observer Z.1 inverted microscope with a Photometrics Evolve EMCCD camera. ZEN 2.6 software was used for acquisition. Tiles and Z-series were stitched using ZEN software for whole larvae images.

### Imaging analyses and processing

Images were processed and analyzed using FIJI ImageJ (62). Maximum intensity projections of Z-series images were used for representative images and further analysis of the tailfin and leukocyte area. For measuring tissue regrowth, the total fin tissue area distal to the notochord was outlined using the polygon tool (41). Leukocyte recruitment and bacterial area were analyzed by manual thresholding and measuring the 2D area of the corresponding fluorescent signal as depicted in Figure 2A-B. In all representative images, brightness and contrast were adjusted for visual purposes only, and no alterations were made to images before analysis. Three biological replicates were run for all experiments, and the total number of fish used is listed in the appropriate figure legends. Once data was collected, distribution normality was assessed with the use of a D’Agostino-Pearson omnibus normality test, and statistical significance was determined via either Student’s t-test, multiple Student’s t-tests, or multiple Mann–Whitney tests. RT-qPCR data statistical significance was determined by Two-way ANOVA (GraphPad Prism, v7.0c software).

## Acknowledgements

We thank Dr. John-Demian Sauer (University of Wisconsin-Madison) for his generosity in providing the *L. monocytogenes* strain used in this study. Schematics in Figures 1, 4, Supplementary Figure 1 and 3 were created using BioRender.com

This work was supported by the National Science Foundation (NSF) Graduate Research Fellowship Program under Grant No. (DGE-2137424) awarded to N.M.S., and R35 GM118027 to AH. Any opinions, findings, and conclusions or recommendations expressed in this material are those of the author(s) and do not necessarily reflect the views of the NSF or the NIH.

---

**Supplemental Figure 1.**
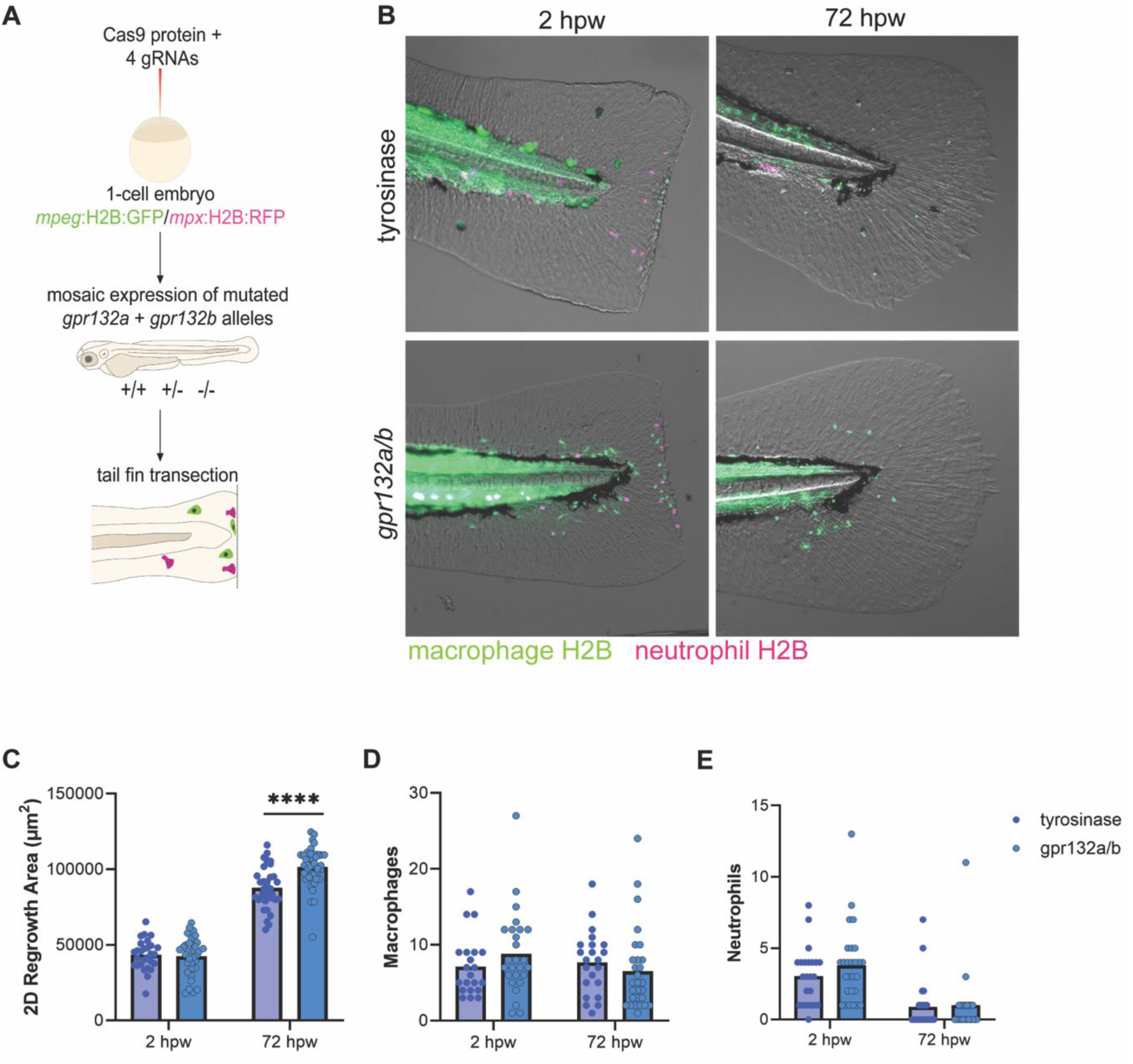
Gpr132a/b crispants heal better upon tail transection. (A) Schematic for the generation of *gpr132a/b* crispant larvae. *mpeg*:H2B:GFP/*mpx*:H2B:RFP embryos were injected with Cas9 and gRNAs at single-cell stage. Larvae were wounded at 3 dpf and fixed at 2 and 72 hptt. (B) Representative tail fin images of double transgenic *mpeg*:H2B:GFP/*mpx*:H2B:RFP (green macrophages and red neutrophils) *gpr132a/b* crispant larvae at 2 and *72* hptt. Control larvae were injected with gRNAs targeting tyrosinase. (C) 2D regrowth area of *gpr132a/b* crispants vs control injected larvae at 2 and 72 hptt. (D) Number of macrophages (*mpeg*:H2B:GFP) of *gpr132a/b* crispants vs control injected larvae at 2 and 72 hptt. (E) Number of neutrophils (*mpx*:H2B:RFP) of *gpr132a/b* crispants vs control injected larvae at 2 and 72 hptt. Bars on graphs are presented as mean + standard deviation. Each dot represents data from an individual fish, and results represent data pooled from 3 independent experiments. n = 24-30 larvae per condition. *P*-values calculated by Multiple Mann-Whitney tests. \*\*\*\**p*<0.0001.

**Supplemental Figure 2.**
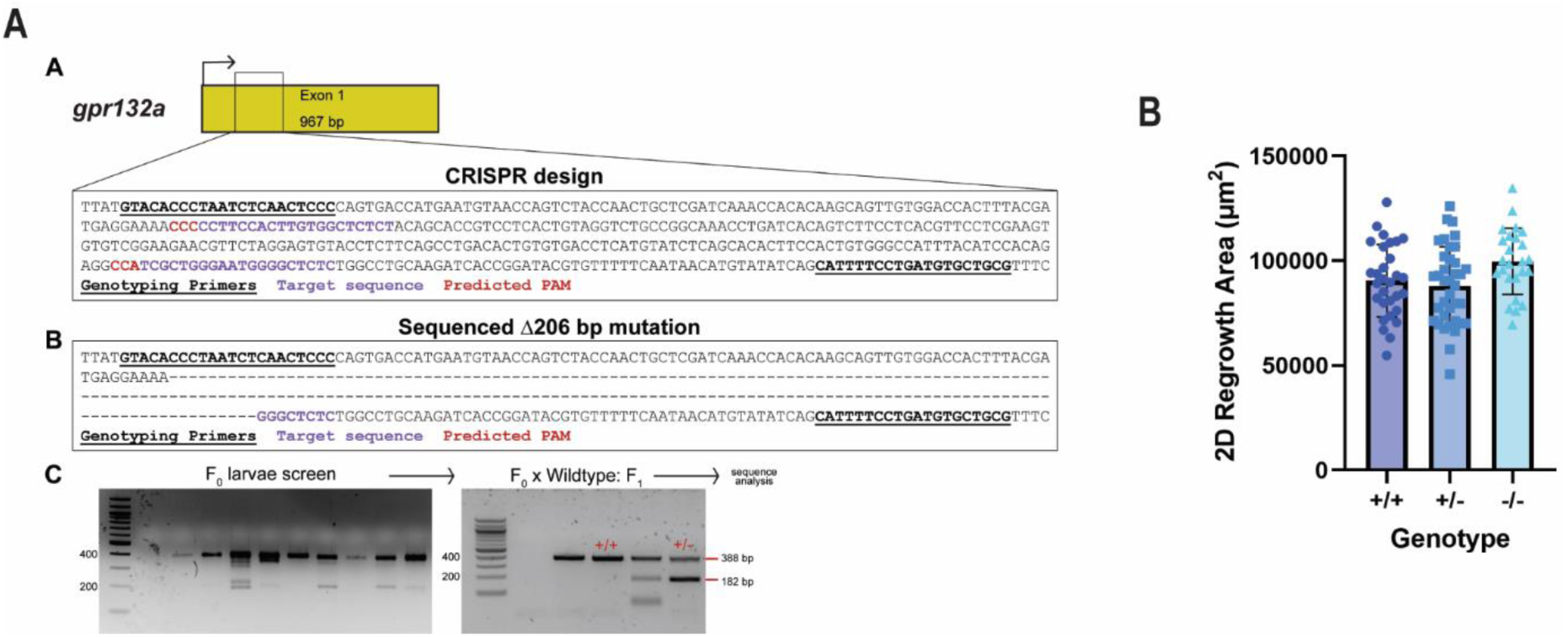
Gpr132a does not influence wound repair. (A) Schematic of *gpr132a* gene including the target region for CRISPR/Cas9 mutagenesis. Two guide RNAs (gRNAs) were used for gene editing. Genotyping primers are indicated with bold underlined text, gRNA target sequences are indicated in purple, and the PAM sequences predicted by CRISPRscan.org are indicated in red. Cutting at both gRNA target sequences generated a 206 bp deletion in exon 1 as determined by sequencing. Genotyping of F0 larvae followed by sequential outcrossing of heterozygous mutants containing the 206 bp deletion. (B) 2D Regrowth area at 72 hptt of a *gpr132a* heterozygous incross, including all genotypes. Bars on graphs are presented as mean + standard deviation. Each dot represents data from an individual fish, and results represent data pooled from 3 independent experiments. n = 24-30 larvae per condition. *P*-values calculated by Two-way ANOVA with Tukey’s multiple comparisons tests.

**Supplemental Figure 3.**
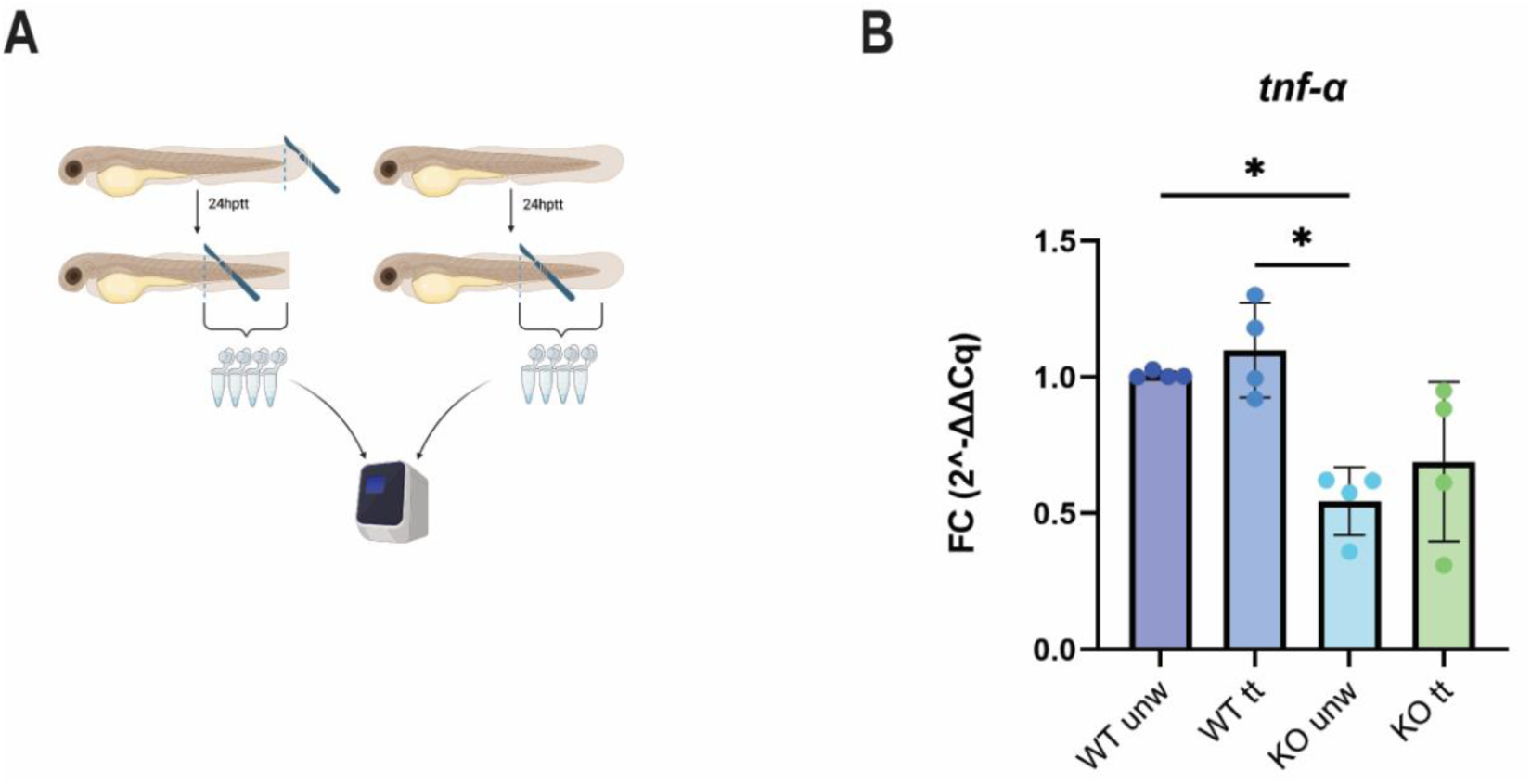
Gpr132b-deficient larvae exhibit lower basal *tnf-a* expression. (A) Larvae from each genotype were either transected (∼50) or left unwounded (∼50) at 3 dpf. After 24 hours, tissue from the urogenital opening to the tail fin was dissected and pooled for RNA extraction, cDNA synthesis, and qPCR. (B) RT-qPCR of *tnf-a* expression in posterior tissue at 24 hptt or 4dpf. Dots represent fold change in gene expression from pools of ∼50 larvae. Results represent 4 independent experiments. Fold change was calculated using the ddCq method. P values calculated with Two-way ANOVA. \**p*<0.05.

